# Genomic features and instability of linezolid resistance in vancomycin-resistant *Enterococcus faecalis* ST525

**DOI:** 10.64898/2026.11.04.764223

**Authors:** Lara Mendes de Almeida, Josiane Trevisol Leal, Laís Carolina Scapolan Ito, Kawany Miyazaki Araujo, Felipe Mendes Zorzi, Joas Lucas Silva, Andrea Micke Moreno, Ana Cristina Gales

## Abstract

This study investigated the mechanisms of oxazolidinone resistance and genomic background in clinical ST525 *Enterococcus faecalis* isolates, a poorly characterized lineage reported in Brazilian hospitals as either vancomycin-resistant (VRE) or linezolid- and vancomycin-resistant (LVRE). Five LVREfs from intensive care unit patients (linezolid MICs, 8–32 mg/L; vancomycin MICs, >256 mg/L) carried only G2576T and G2505A (in three of four *rrl* copies) as known linezolid resistance determinants, despite variable linezolid MICs. During serial *in vitro* passages without antibiotic pressure, isolates with initial MICs of 8–16 mg/L exhibited progressive MIC reductions accompanied by reversion of mutant to wild-type *rrl* copies, whereas the isolate with an initial MIC of 32 mg/L maintained high-level resistance without reversion. Additional domain V substitutions emerged during MIC decline, while growth kinetics remained broadly similar. Whole-genome sequencing of a representative isolate (LVREfs L7) revealed a ∼2.8-Mb chromosome enriched in mobile genetic elements (∼8.6%), lacking a functional CRISPR–Cas system. A plasmid-borne prototype Tn*1546* carrying *vanA*, devoid of internal insertion sequences, was found inserted into a Tn*3*-family transposase gene, a platform not typically associated with *vanA* mobilization. Together, these findings indicate that ST525 LVREfs, whose genomic features are consistent with long-term persistence in hospitals, may pose a challenge to routine susceptibility testing. A single mutational mechanism can yield variable linezolid MICs, and the ratio of mutated vs wild-type *rrl* copies can become heterogeneous in the absence of antibiotic pressure, without correlating directly with these values, contrary to the well-documented copy number–MIC correlation.

## Introduction

Vancomycin-resistant enterococci (VRE) are a major cause of healthcare-associated infections worldwide. In Brazil, initial reports date from the late 1990s ^1^, approximately a decade after their emergence in the United States and Europe. Shortly thereafter, VRE incidence increased substantially, rising from 4% in 2003 to 12% in 2008 ^2^. By 2011, the country accounted for the highest proportion of VRE cases in Latin America, reaching 27% ^3^. More recent nationwide surveillance data (2019–2024) confirm an upward trend in vancomycin resistance among *Enterococcus* isolates. VRE rates rose sharply during the COVID-19 pandemic, and although they have declined somewhat in the post-pandemic period, they remain high, with *Enterococcus faecium*–associated laboratory-confirmed bloodstream infections (BSIs) and urinary tract infections (UTIs) now exceeding 50% in adult and pediatric intensive care units (ICUs) ^4^.

The introduction of the first oxazolidinone, linezolid, provided an important therapeutic option against VRE and other multidrug-resistant (MDR) Gram-positive pathogens, including methicillin-resistant *Staphylococcus aureus* (MRSA), methicillin-resistant coagulase-negative staphylococci (MRSCoN), and penicillin-resistant *Streptococcus pneumoniae* (PRSP) ^5^. Linezolid has also become a drug of choice in treatment regimens for MDR *Mycobacterium tuberculosis* and MDR nontuberculous mycobacteria infections ^6^. However, continuous and intensive use of antibiotics in both hospital and agricultural settings has intensified bacterial adaptation through both mutational and horizontally acquired oxazolidinone resistance mechanisms ^7^, further exacerbating the already critical challenge posed by isolates nonsusceptible to glycopeptides and β-lactams.

Despite the contribution of the *cfr*, *optrA*, and *poxtA* genes to oxazolidinone resistance, mutations in the central loop of domain V of 23S rRNA (*rrl*), approximately 6–8 Å from the linezolid-binding site, continues to pose a major challenge to linezolid therapy. These mutations reduce linezolid binding affinity within the peptidyl transferase center (PTC) of the 50S ribosomal subunit, which prevents inhibition of aminoacyl-tRNA accommodation at the A-site and avoids disruption of fMet-tRNA positioning at the P-site, thereby allowing proper initiation complex formation during translation ^8^. Multiple mutated *rrl* copies, particularly those harboring the G→T transversion at nucleotide 2576, have consistently been associated with linezolid minimum inhibitory concentrations (MICs) exceeding the Clinical and Laboratory Standards Institute (CLSI) and European Committee on Antimicrobial Susceptibility Testing (EUCAST) breakpoints in clinical isolates. Although both clinical discontinuation of linezolid therapy and *in vitro* serial passage in antibiotic-free media have been shown to restore wild-type *rrl* copies and reduce MICs ^9–13^, interpretation can be challenging because MIC shifts toward clinical breakpoints complicate genotype-phenotype correlations, as recently demonstrated in a multicenter study on enterococci with borderline MICs of 4–8 mg/L ^14^. Moreover, conventional resistance detection methods, including automated identification and susceptibility testing systems routinely employed in hospital laboratories, cannot distinguish isolates harboring different numbers of mutated *rrl* copies and may consequently classify some of them as linezolid susceptible despite the presence of resistance-associated mutations. Even when these mutations are ruled out before treatment, a screening rarely performed in routine clinical practice, preferential transcription of mutated *rrl* over wild-type copies during therapy can lead to inconsistent results across susceptibility testing methods, ultimately contributing to therapeutic failures.

Given the reliance on oxazolidinone-based therapies for treating VRE and other MDR Gram-positive infections, a comprehensive understanding of emerging lineages resistant to these compounds is essential to strengthen surveillance and inform clinical decision-making. Therefore, we investigated the genetic basis underlying oxazolidinone resistance and genomic background of linezolid- and vancomycin-resistant *Enterococcus faecalis* (LVREfs) isolates previously shown to harbor the G2576T mutation in domain V of 23S rRNA and the glycopeptide resistance gene *vanA* ^15^. Our renewed interest in these isolates (linezolid MICs, 8–32 mg/L; vancomycin MIC, 256 mg/L) stems from their assignment to sequence type (ST) 525, a poorly characterized lineage that appears to be endemic in Brazil ^15–17^. We investigated whether oxazolidinone resistance could be mediated by mechanisms described after these LVREfs were initially reported ^15^ and evaluated the stability of their mutated *rrl* copies. In addition, we characterized both chromosomally integrated and plasmid-borne mobile genetic elements (MGEs) and assessed whether deficiencies in clustered regularly interspaced short palindromic repeat (CRISPR) systems may have facilitated their acquisition.

## Results

### Heterogeneity of *rrl* in ST525 LVREfs during antibiotic-free serial passage

We screened all ST525 LVREfs isolates for known oxazolidinone resistance determinants and monitored their stability during 100 days of serial passage in antibiotic-free medium. We also compared the growth of the parental isolates with that of their derivatives showing reduced linezolid MICs. All encoded wild-type ribosomal proteins L3, L4, and L22 and lacked *cfr*, *optrA* and *poxtA*. At day 1 of serial passages, three of the four *rrl* copies in each isolate carried both G2576T and G2505A within domain V (**Table 1**), while linezolid MICs ranged from 8 to 32 mg/L. During daily subculturing, isolates with initial linezolid MICs of 8 or 16 mg/L showed progressive reductions in MIC values, concomitant with the restoration of wild-type *rrl* copies. The isolate with an initial linezolid MIC of 32 mg/L retained its high-level resistance and three mutated *rrl* copies throughout the 100-day serial passage assay (**Figure 1**). Four additional substitutions emerged in domain V as linezolid MICs declined: G21A and C1468T were identified in *rrl2* and *rrl3*, respectively, after G2576T and G2505A were no longer detected in those copies, while G401A and T2305G were detected in *rrl4* in individual isolates (**Table 1**). Growth curves were broadly similar between the parental LVREfs isolates and their corresponding serial-passage derivatives with reduced linezolid MICs.

**Figure 1.**
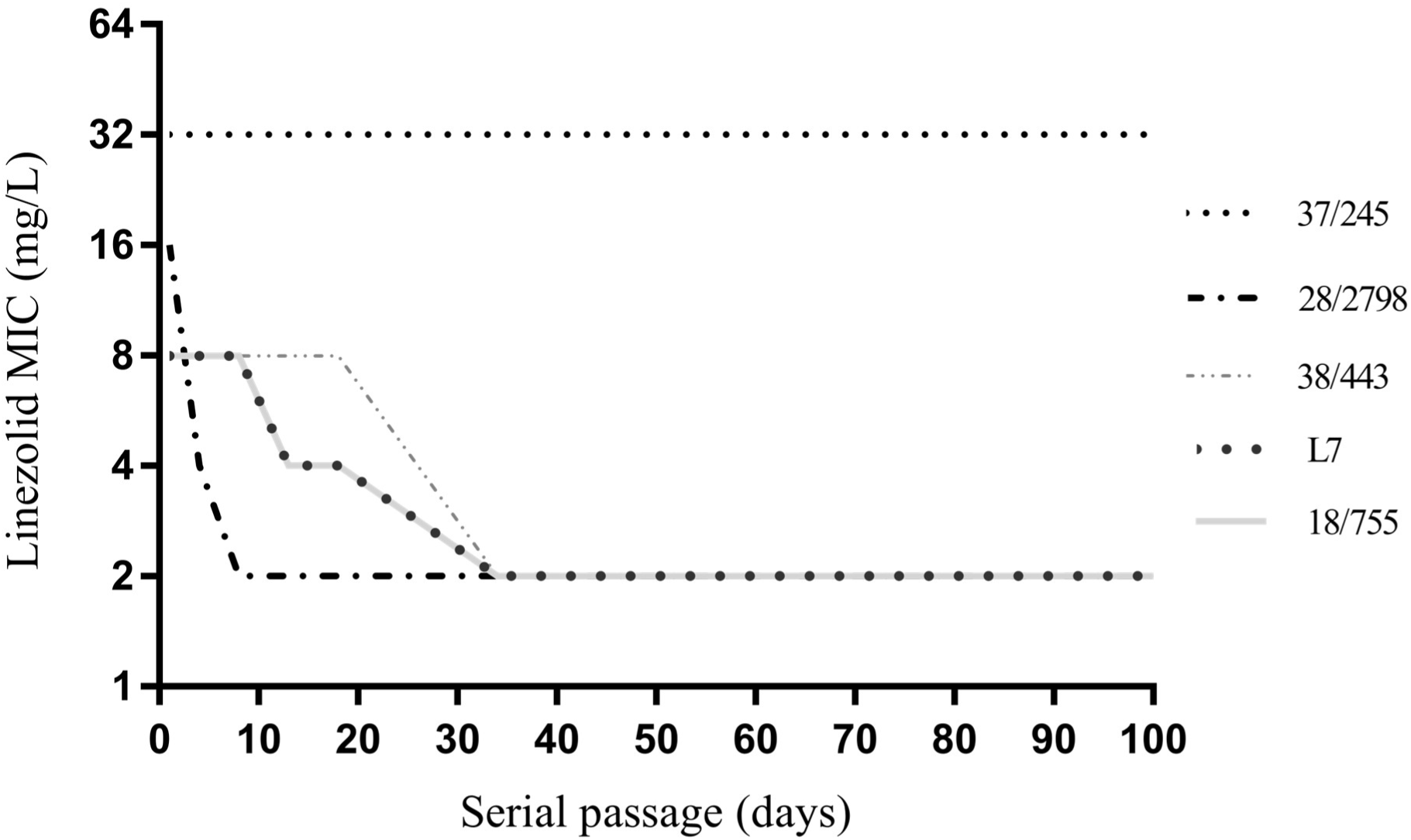
Linezolid minimum inhibitory concentrations (MICs) over 100 days of serial subculturing in ST525 linezolid- and vancomycin-resistant *Enterococcus faecalis* (LVREfs) isolates. Reduction and stability of linezolid MICs are shown for isolates L7, 18/755, 38/443, 28/2798, and 37/245 according to Clinical and Laboratory Standards Institute (CLSI) M100 breakpoints: susceptible, ≤2 mg/L; intermediate, 4 mg/L; resistant, ≥8 mg/L.

**Table 1.** Domain V rrl substitutions in ST525 LVREfs isolates during serial subculturing.

| Isolate | copy | Day 1 | Day 4 | Day 8 | Day 13 | Day 18 | Day 34 | Day 100 |
|---|---|---|---|---|---|---|---|---|
| L7 | *rrl1* | wt | wt | wt | wt | wt | wt | wt |
|  | *rrl2* | G2576T<br>G2505A | G2576T<br>G2505A | G2576T<br>G2505A | G2576T<br>G2505A | G21A | G21A | G21A |
|  | *rrl3* | G2576T<br>G2505A | G2576T<br>G2505A | G2576T<br>G2505A | G2576T<br>G2505A | G2576T<br>G2505A | C1468T | C1468T |
|  | *rrl4* | G2576T<br>G2505A | G2576T<br>G2505A | G2576T<br>G2505A | G2576T<br>G2505A<br>G401A | G2576T<br>G2505A<br>G401A | G2576T<br>G2505A<br>G401A | G2576T<br>G2505A<br>G401A |
| 18/755 | *rrl1* | wt | wt | wt | wt | wt | wt | wt |
|  | *rrl2* | G2576T<br>G2505A | G2576T<br>G2505A | G2576T<br>G2505A | G2576T<br>G2505A | G21A | G21A | G21A |
|  | *rrl3* | G2576T<br>G2505A | G2576T<br>G2505A | G2576T<br>G2505A | G2576T<br>G2505A | G2576T<br>G2505A | C1468T | C1468T |
|  | *rrl4* | G2576T<br>G2505A | G2576T<br>G2505A | G2576T<br>G2505A | G2576T<br>G2505A<br>G401A | G2576T<br>G2505A<br>G401A | G2576T<br>G2505A<br>G401A | G2576T<br>G2505A<br>G401A |
| 38/443 | *rrl1* | wt | wt | wt | wt | wt | wt | wt |
|  | *rrl2* | G2576T<br>G2505A | G2576T<br>G2505A | G2576T<br>G2505A | G2576T<br>G2505A | G21A | G21A | G21A |
|  | *rrl3* | G2576T<br>G2505A | G2576T<br>G2505A | G2576T<br>G2505A | G2576T<br>G2505A | C1468T | C1468T | C1468T |
|  | *rrl4* | G2576T<br>G2505A | G2576T<br>G2505A | G2576T<br>G2505A | G2576T<br>G2505A | G2576T<br>G2505A<br>T2305G | G2576T<br>G2505A<br>T2305G | G2576T<br>G2505A<br>T2305G |
| 28/2798 | *rrl1* | wt | wt | wt | wt | wt | wt | wt |
|  | *rrl2* | G2576T<br>G2505A | wt | G21A | G21A | G21A | G21A | G21A |
|  | *rrl3* | G2576T<br>G2505A | G2576T<br>G2505A | G2576T<br>G2505A | wt | wt | wt | wt |
|  | *rrl4* | G2576T<br>G2505A | G2576T<br>G2505A | G2576T<br>G2505A | G2576T<br>G2505A | G2576T<br>G2505A | G2576T<br>G2505A | G2576T<br>G2505A |
| 37/245 | *rrl1* | wt | wt | wt | wt | wt | wt | wt |
|  | *rrl2* | G2576T<br>G2505A | G2576T<br>G2505A | G2576T<br>G2505A | G2576T<br>G2505A | G2576T<br>G2505A | G2576T<br>G2505A | G2576T<br>G2505A |
|  | *rrl3* | G2576T<br>G2505A | G2576T<br>G2505A | G2576T<br>G2505A | G2576T<br>G2505A | G2576T<br>G2505A | G2576T<br>G2505A | G2576T<br>G2505A |
|  | *rrl4* | G2576T<br>G2505A | G2576T<br>G2505A | G2576T<br>G2505A | G2576T<br>G2505A | G2576T<br>G2505A | G2576T<br>G2505A | G2576T<br>G2505A |

Only time points at which at least one isolate showed a reduction in linezolid MIC during antibiotic-free serial passage are included; wt, wild type.

### Clonal relatedness and genomic overview of LVREfs L7

LVREfs L7 was selected for whole-genome sequencing (WGS) as a representative of the predominant *Sma*I pulsed-field gel electrophoresis (PFGE) pulsotype, shared by three of the five ST525 isolates previously recovered from BSIs and UTIs ^15^. Genome assemblies of L7 yielded a ∼2.8-Mb chromosome, of which approximately 8.6% corresponded to MGEs. These included four intact prophages (36.8–46.2 kb), a 17,990-bp T4SS-type Tn*916*-like integrative and conjugative element (ICE) carrying *tet*(M), and a 5,782-bp integrative mobilizable element (IME) with *dfrF*. Additional chromosomal features comprised *lsa*(A), genes conferring biocide and metal tolerance, alongside virulence-associated determinants (*ace*, *efaAfs*, *ebpABC*, *agg*, *fsrB*, *gelE*, *hylA*, and *hylB*). Although plasmid structures were only partially resolved by short-read assemby, the inferred plasmidome included RepA_N-type and Inc18 replicons and a MobP relaxase. Putative plasmid-associated resistance determinants included *cat*, *erm*(B), and a complete *vanA*-carrying Tn*1546*. The 10.8-kb Tn*1546* closely matched the prototype sequence from *E. faecium* BM4147 (GenBank accession no. M97297), containing *vanR*, *vanS*, *vanH*, *vanA*, *vanX*, *vanY*, and *vanZ* ^18^. Disrupted fragments (1,161 bp and 1,854 bp) of a Tn*3*-family transposase gene (NZ_CABHGG010000009) flanked *orf1* and *vanZ*, respectively. BLASTn analysis of the complete 13.8-kb region returned no full-length matches. As for CRISPR-Cas defense, only a short orphan CRISPR2 array containing two spacers was identified, with no associated *cas* genes. Both spacers showed 100% identity and coverage in BLASTn searches: spacer 1 returned approximately 100 matches, including a *Caudoviricetes* sequence recovered from an urban metagenome ^19^, whereas spacer 2 matched approximately 15 sequences from hospital-associated *E. faecalis* strains.

## Discussion

Reports of ST525 *E. faecalis* in southeastern Brazil coincided with a shift in the epidemiology of *vanA*-positive VRE from *E. faecalis* to *E. faecium* predominance beginning in the mid-2000s ^20^. The limited number of isolates and single-center sampling do not establish ST525 as a broadly disseminated or hospital-adapted lineage; however, its persistence during this epidemiological transition and over a three-year interval in the same hospital is consistent with its capacity for long-term persistence in clinical settings. Its origin and evolutionary trajectory remain uncertain; nevertheless, available pubMLST data place ST525 *E. faecalis* at a peripheral position relative to hospital-adapted *E. faecalis* lineages and suggest it may have evolved from the predicted founder of CC2 *E. faecalis* ST6 through intermediate sequence types, potentially including ST738 or ST1770.

Direct correlations between linezolid MICs and the proportion of *rrl* copies harboring G2576T or G2505A have been previously demonstrated, with mutated copy counts increasing under linezolid exposure and declining, alongside MIC reductions, due to wild-type *rrl* restoration upon drug withdrawal ^9–13^. More recently, a study investigating how *E. faecalis* adapts to increasing linezolid pressure through distinct, concentration-dependent strategies revealed that G2505A emerges at higher concentrations to consolidate resistance despite imposing a fitness cost^21^. In the small set of isolates analyzed here, the presence of G2576T and G2505A in three of four *rrl* copies did not translate into the classic stepwise change in MICs linked to the dosage of mutated *rrl* copies, suggesting other underlying factors. Additional functional studies are required to determine the role of substitutions that emerged as linezolid MICs declined (G21A, G401A, C1468T, and T2305G). The broadly similar growth curves do not establish whether they alter competitive fitness or serve as an adaptive strategy to preserve essential 23S rRNA functions while offsetting potential fitness costs imposed by G2576T and G2505A.

The repertoire of MGEs in *Enterococcus* genomes provides valuable insight into their recent evolutionary trajectories and ecological interactions. Hospital-adapted MDR *Enterococcus* lineages typically possess expanded genomes due to MGE accumulation, often ∼25% larger than those of antibiotic-susceptible commensal isolates, a feature enabled by reduced CRISPR-Cas-mediated restriction, given that a functional system limits plasmid and phage uptake ^22,23^. While the LVREfs L7 genome lacks the expanded size typical of entrenched MDR hospital lineages, its mobilome suggests that it may be evolving in this direction rather than reflecting passive MGE acquisition, consistent with its CRISPR-deficient genome. Although its CRISPR2 array may serve alternative functions, the limited spacer content suggests minimal exposure to MGEs, which is consistent with a relatively recent introduction into the hospital environment. In line with this, L7 Tn*1546* showed a non-mosaic configuration, in contrast to the insertion-sequence-rich structures commonly reported in *E. faecium* in Brazil ^24,25^, suggesting infrequent chromosomal or interplasmid recombination events. Interestingly, the Tn*3*-family platform into which this element was inserted may provide an alternative route for *vanA* mobilization.

## Conclusions

In this small isolate set, G2576T and G2505A yielded variable linezolid MICs despite an equivalent initial *rrl* copy distribution. The loss of linezolid resistance during antibiotic-free passage appeared to be associated with heterogeneous changes in *rrl* copy composition, including additional substitutions, rather than with a simple reduction in the number of mutated copies. Further investigations should clarify this heterogeneity and the persistence of high-level resistance in the isolate with an MIC of 32 mg/L, which resisted wild-type *rrl* restoration even in the absence of selective pressure. Such reversible heterogeneity among *rrl* copies may obscure the interpretation of linezolid susceptibility phenotypes, potentially allowing resistance-associated mutations to escape routine detection and increasing the risk of oxazolidinone therapy failure. The genomic features of LVREfs L7 may not be shared by all ST525 *E. faecalis*, but they are consistent with long-term persistence in the hospital environment. Together with the atypical flanking regions of its Tn*1546*, they underscore the need for continuous genomic surveillance for the emergence of new VRE and LVRE populations.

## Materials and Methods

### Assessment of persistence of domain V *rrl* mutations and other oxazolidinone resistance determinants

Linezolid resistance-associated domain V mutations in *rrl* copies were identified using nested PCR. In the first-round PCR, a single forward primer annealing to a conserved region approximately 100 bp upstream of domain V was combined with four reverse primers targeting the variable 16S–23S intergenic spacer (ITS1) regions ^13^. Outer amplicons were used as templates for the second-round PCR, which employed inner primers annealing to the terminal regions of domain V. Nested PCR products were subjected to Sanger sequencing and aligned to the corresponding 23S rRNA domain V sequences of linezolid-susceptible *E. faecalis* reference strains, with nucleotide positions numbered according to the *Escherichia coli* 23S rRNA sequence. Persistence of mutations in individual *rrl* copies was assessed by nested PCR at each reduction in linezolid MIC values during serial subculture of LVREfs isolates on antibiotic-free Mueller– Hinton medium. MICs for linezolid, tedizolid, and vancomycin were determined by broth microdilution, according to ISO 20776-1 ^26^.

The impact of domain V *rrl* mutations on bacterial fitness was evaluated by comparing growth curves of LVREfs isolates at day 1 with those of their corresponding serial-passage derivatives exhibiting reduced linezolid MICs ^27^. Overnight cultures were diluted in fresh medium and adjusted to an OD₆₀₀ of 0.08–0.10 (≈10⁸ CFU/mL). Cultures were inoculated at a 1:20 ratio into 200 μL of tryptic soy broth (TSB) in 96-well microplates and incubated at 37°C for 24 h. Optical density at 600 nm was measured hourly using a BioTek microplate reader. Growth kinetics were determined from OD₆₀₀ values over time, and data represent the mean of quadruplicate experiments.

Mutations in the ribosomal uL3, uL4, and uL22 proteins were assessed by PCR of their encoding *rplC*, *rplD*, and *rplV* genes, respectively, using primers annealing to the terminal regions of each gene, followed by Sanger sequencing of the resulting amplicons. The *cfr*, *optrA* and *poxtA* genes were investigated by PCR as previously described ^28^.

### Genome sequencing and analysis

As representative of the predominant PFGE pulsotype, genomic DNA of isolate LVREfs L7 was extracted using the DNeasy Blood & Tissue Kit (Qiagen, Germantown, MD, USA) and sequenced on an Illumina MiSeq platform, generating 2 × 250 bp paired-end reads from libraries prepared with the Nextera XT DNA library preparation kit (Illumina). Raw reads were assessed for quality using FastQC v0.11.9 ^29^ and Centrifuge v1.0.4 ^30^, and filtered with Fastp v0.23.4 ^31^. For comparative evaluation, assemblies *de novo* using Velvet (via PLACNETw) ^32^, SPAdes 3.15.5 ^33^, Unicycler v0.5.0 ^34^, and Shovill v1.1.0 ^35^ were performed. Assembly quality was assessed using QUAST v5.2.0 ^36^ and CheckM v1.2.3 ^37^, and chromosomal scaffolding was generated with RagTag v2.1.0 ^38^. For genome annotation, the Rapid Annotation using Subsystem Technology (RAST) server ^39^, Bakta v1.9.4 ^40^, and the NCBI Prokaryotic Genome Annotation Pipeline (PGAP) ^41^ were used. Analyses employed BLAST+ tools ^42^, Geneious R9.0.5 utilities <https://www.geneious.com/>, and resources from the Center for Genomic Epidemiology <https://www.genomicepidemiology.org/>, as well as CRISPRCasFinder v.4.3.2 ^43^, PHASTER v.2.0 ^44^, oriTFinder2 v.2.0 ^45^, and ICEberg 3.0 ^46^.

## Data availability

Whole-genome sequencing data have been deposited in the DDBJ/ENA/GenBank database under the accession number JBTOFM000000000.

## Acknowledgements

We gratefully acknowledge Prof. Dr. Elsa Masae Mamizuka (Department of Clinical and Toxicological Analyses, University of São Paulo, SP, Brazil) for providing laboratory facilities for part of this work.

## Author contributions

Lara M. Almeida: conceptualization, investigation, data curation, formal analysis, and writing– original draft, review and editing; Josiane Trevisol Leal: investigation, data curation, formal analysis, and writing– original draft, review and editing; Laís Carolina Scapolan Ito: investigation; Kawany M. Araujo: formal analysis and software; Felipe M. Zorzi: formal analysis and software; Joas L. Silva: formal analysis and writing– review and editing; Andrea Micke Moreno: writing– review and editing, and funding acquisition; Ana Cristina Gales: conceptualization, supervision, writing– review and editing, and funding acquisition.

## Declarations

### Ethical approval

This study used bacterial isolates from a collection previously held by a laboratory at the University of São Paulo (USP) and did not involve human participants or identifiable patient data. Therefore, ethical approval and informed consent were not required.

